# Hippocampal Spine Head Sizes Are Highly Precise

**DOI:** 10.1101/016329

**Authors:** Thomas M Bartol, Cailey Bromer, Justin Kinney, Michael A. Chirillo, Jennifer N. Bourne, Kristen M. Harris, Terrence J Sejnowski

**Affiliations:** Howard Hughes Medical Institute, the Salk Institute for Biological Studies, La Jolla, CA 92037; McGovern Institute for Brain Research, Massachusetts Institute of Technology, Cambridge, MA, 02139; Center for Learning and Memory, Department of Neuroscience, University of Texas, Austin, TX 78712-0805, USA; Division of Biological Sciences, University of California at San Diego, La Jolla, CA 92093

**Author notes:** J. Kinney - Current address: MIT. J. Bourne - current address: Univ. Colorado, Denver.

## Abstract

Hippocampal synaptic activity is probabilistic and because synaptic plasticity depends on its history, the amount of information that can be stored at a synapse is limited. The strong correlation between the size and efficacy of a synapse allowed us to estimate the precision of synaptic plasticity. In an electron microscopic reconstruction of hippocampal neuropil we found single axons making two or more synaptic contacts onto the same dendrites which would have shared histories of presynaptic and postsynaptic activity. The postsynaptic spine heads, but not the spine necks, of these pairs were nearly identical in size. The precision is much greater than previous estimates and requires postsynaptic averaging over a time window many seconds to minutes in duration depending on the rate of input spikes and probability of release.

**One Sentence Summary:** Spine heads on the same dendrite that receive input from the same axon are the same size.

---

Excitatory synapses on dendritic spines of hippocampal pyramidal neurons have a wide range of sizes that are highly correlated with their synapse sizes and strengths ^1–7^. Due to high failure rate and other sources of stochastic variability, these synapses transmit unreliably individual presynaptic action potentials. Nonetheless, the sizes and strengths of these synapses can increase or decrease according to the relative timing of presynaptic inputs and postsynaptic spikes ^8^. Prior work suggests that pairs of spines on the same dendrite contacting the same axon are more similar in size than those from the same axon on different dendrites ^9^. Here we evaluated this axon-spine coupling in three-dimensional reconstructions from serial electron microscopy reconstruction (3DEM) of hippocampal neuropil from 3 adult rats, to determine this similarity with higher precision and the time window over which pre- and post-synaptic histories would need to be coordinated.

In a 6 x 6 x 5 µm^3^ complete 3DEM from the middle of stratum radiatum in hippocampal area CA1 ^10,11^, we identified 149 dendritic branches, 446 axons, and 449 synapses. We measured spine head volume and surface area, surface area of the postsynaptic density (PSD) adjacent to the presynaptic active zone, and quantified the number of vesicles at the 288 spines fully contained within the volume. Strong correlations between these metrics are consistent with previous observations ^1,4^ and confirm the typicality of our sample (Figs. 1, 2, S1). To reduce error, we averaged over multiple independent spine volume measurements for each spine (Fig. 1A, Fig. S2).We determined that the relationship between PSD area and spine head volume did not differ significantly across different dendritic branches (Fig. S3). The correlation between spine head area and spine head volume accounted for 99% of the variance, despite the wide range in spine head shapes and dimensions (Fig. S1E), which suggests that the accuracy of our measurements matched the precision of the spine. We also measured spine neck volume and found no significant correlation between the neck and PSD area (Fig. 1C) or spine head volume (Fig. 1D).

**Fig. 1.**
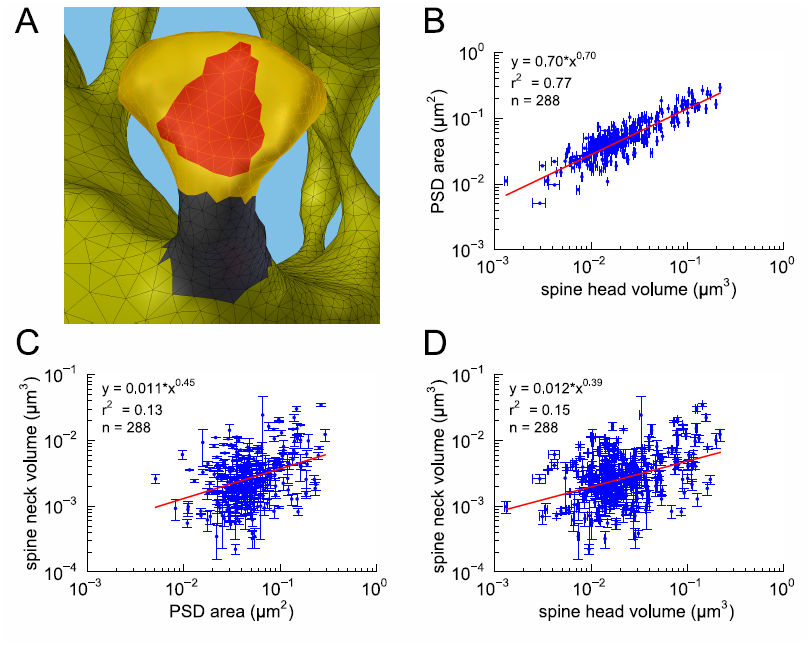
Spine head volumes, but not neck volumes, are correlated with PSD areas. A) Example segmentation of spine head (yellow), neck (gray), and PSD area (red). B) Strong correlation between PSD area and spine head volume. No correlation between spine neck volume and C) PSD area or D) spine head volume. Regression lines in red and error bars for each data point represent SEM based on multiple tracers who also edited each spine. Equations are based on the log-log distributions, with r^2^ values indicated, and n=288 complete spines.

**Fig. 2.**
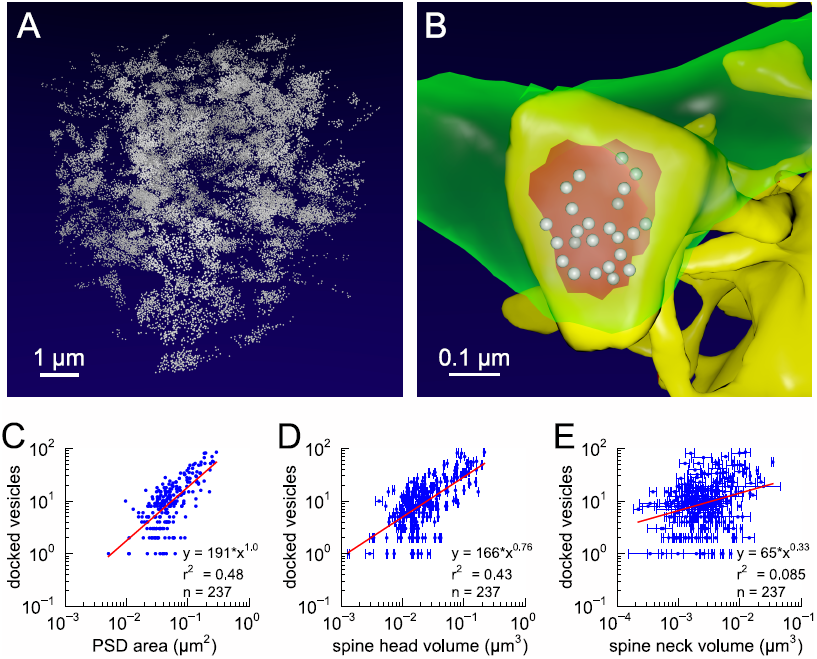
Presynaptic docked vesicle numbers are correlated with PSD areas and spine head volumes, but not with spine neck volumes. A) All 31,377 presynaptic vesicles. B) *En face* view of the 24 docked vesicles (gray spheres) viewed through an axon (green) onto the PSD (red) of example spine (yellow). C) Number of docked vesicles is correlated with PSD area and D) spine head volume, but not correlated with E) spine neck volume. Regression lines, SEM, and r^2^ are as in Fig. 1, n = 237 complete axonal boutons, each associated with one of the 288 complete spines. One tracer marked vesicles, hence no SEM.

Next, we analyzed spine volumes according to their axonal connectivity and dendritic origin. Pairs of spines on the same dendrite that received input from the same axon (“axon-coupled”), were of the same size and had nearly identical head volumes (Fig. 3, S4-S6). We compared this sample of 10 axon-coupled pairs to those identified from the two additional animals, for a total of 17 pairs. When plotted against one another, the paired head volumes were highly correlated with slope 0.91, and despite the small sample size, were highly significantly different from random pairings of spines (Fig. 3C, S7A, KS, p = 0.0002). Similarly, there was a strong positive correlation between their paired PSD areas (Fig. 3D) and number of presynaptic docked vesicles (Fig. 3E). These features of axon-coupled spines from the same dendrite spanned the distribution of the overall spine population (Fig. S1). In contrast, the spine neck volumes of the pairs were not well-correlated (Fig. 3F), indicating a different function.

**Fig. 3.**
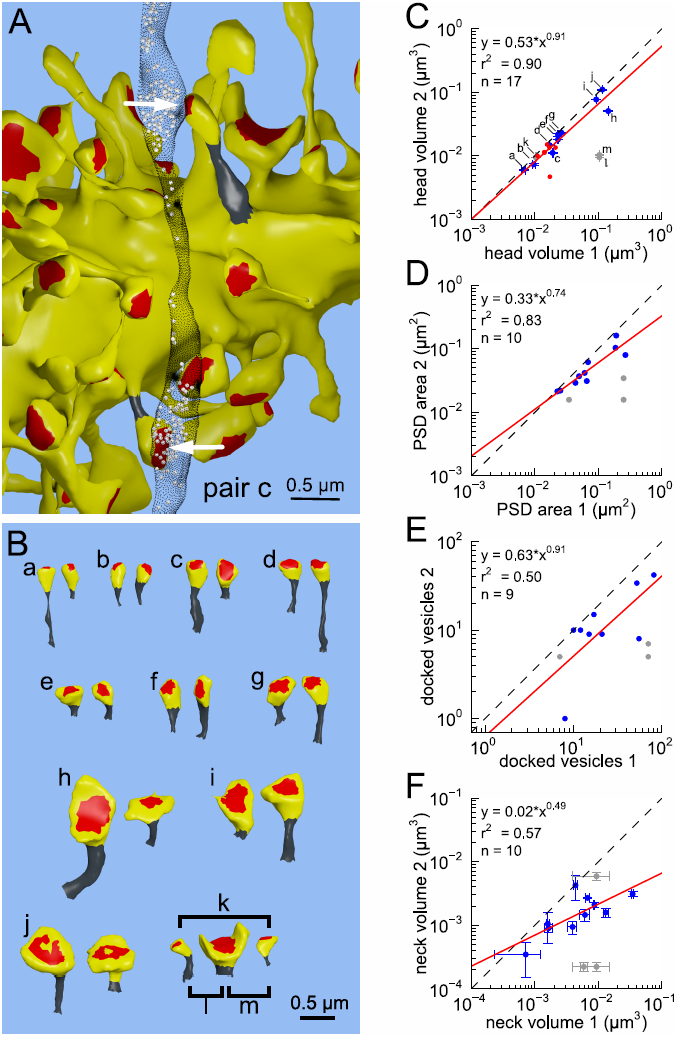
Spine head volumes and PSD areas, but not neck volumes, are highly correlated between pairs of axon-coupled same-dendrite spines. A) Visualization of a pair of spines (gray necks) from the same dendrite (yellow) with synapses (red, indicated by white arrows) on the same axon (black stippling) with presynaptic vesicles (white spheres). B) All axon-coupled same-dendrite spine pairs (colors as in 1A, pair c is elaborated in 3A). Strong correlations with slopes near 1 (dashed diagonal line) occur between paired C) spine head volumes (slope=0.91), D) PSD areas (slope=0.74), and E) docked vesicles (slope=0.91); but not F) spine neck volumes (slope=0.49). Larger values from each pairing are plotted on the X axis. Regression lines (red) include the 10 a-j pairings (blue points) and 7 pairs from 2 additional animals (red points in C), but do not include triplet bouton pairings (k-m, gray points).

The outliers in this set of pairs (Fig. 3C, gray points “k, l, m”) are from comparison of three spines on a single dendritic branch receiving synaptic input from a common multi-synaptic bouton. A larger central spine between two similar in size (Fig. 3B, “k, l, m”) produces one same size pair (“k”) and two different size pairs (“l”, “m”). This unusual configuration is probably driven by processes that differ from the other pairs ^9,12^. Excluding this triple synapse, the median value of the coefficient of variation of volume differences between pairs was CV = 0.083 and was constant across the range of spine sizes (Fig. S8).

This near-identical size relationship does not hold for axon-coupled spines on different dendritic branches (Fig. 4B, CV = 0.39, n = 128, example Fig. 4A), nor for non-axon-coupled spines on the same or different dendrites (Figs. 4E, 4F, example Fig. 4D), all cases that would have different activation histories. The volumes of axon-coupled different-dendrite spines are no different from the volumes of random pairs when plotted against one another (KS, p = 0.89, Figs. 4B, 4C, S7A) and the distribution of their sizes was no different from the whole population (KS, p = 0.44). The number of docked vesicles for pairs on different dendrites (Fig. S9B) is not different from random pairings (KS, p = 0.15). The sizes of pairs of axon-coupled spines on the same or different dendritic branches is unaffected by separation distance (Fig. S10), proximity of glia processes to the synapses (Fig. S11)^13,14^, or location of mitochondria in the axon ^15^.

**Fig. 4.**
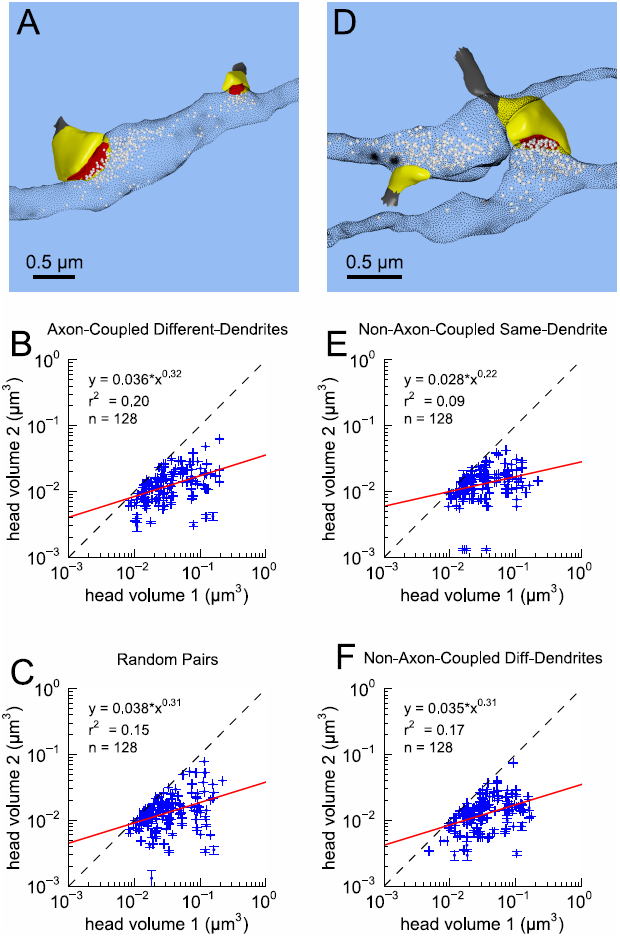
Paired spine head volumes are not correlated when they are not both axon and dendrite coupled. A) Representative visualization and B) plot showing lack of correlation between spine head volumes of all pairs of axon-coupled spines on different dendrites (n=128). C) Similarly, randomly associated pairs of spine head volumes were not correlated. D) Representative visualization and plots show lack of correlation between spine head volumes from randomly selected pairs (n=128) of non-axon-coupled spines E) on the same or F) different dendrites. Color scheme and regression analyses as in Fig. 3.

Spine heads ranged in size over a factor of 60 from smallest to largest, which allows ~24 different strengths to be reliably distinguished across this range, assuming CV = 0.083 and a 75% discrimination threshold (Fig. S12). This corresponds to 4.6 bits of information that can be stored at each synapse (see Methods). The precision of the majority of smaller spines is as good as that of the minority larger spines (Fig. S8), suggesting that accurately maintaining the size of every synapse, regardless of size and strength, could be important for the function, flexibility and computational power of the hippocampus.

How can the high precision in spine head volume be achieved despite the many sources of stochastic variability observed in synaptic responses? These include: 1) The low probability of release from the presynaptic axon in response to an action potential ^5^; 2) Stochastic fluctuations in the opening of postsynaptic NMDA receptors, with only a few of the 2-20 conducting at any time ^16^; 3) Location of release site relative to AMPA receptors ^17–19^ 4) Few voltage-dependent calcium channels (VDCCs) in spines that affect synaptic plasticity (smallest spines contain none) ^20,21^; 5) VDCCs depress after back propagating action potentials ^22^; 6) Capacity for local ribosomal protein synthesis in some spines while others depend on transport of proteins from the dendrites ^23,24^; 7) Homeostatic mechanisms for synaptic scaling may vary ^25,26^; 8) Presence or absence of glia ^13,27^; and 9) Frequency of axonal firing ^28^.

To explain the high precision observed in spine head volumes, we propose that time-window averaging smooths out fluctuations due to plasticity and other sources of variability. To set a lower bound on averaging time we chose to examine neurotransmitter release probability as a single source of variability. Release can be analyzed using a binomial model in which *n* presynaptic action potentials, each with a probability *p_r_* of releasing one or more vesicles, leads to a mean number of releases *µ = n*p*_*r*_ having variance *σ*^2^ = *n*p_r_*(1-p_r_)*. The coefficient of variation around the mean is *CV = sqrt*(*σ*^2^)/µ = *sqrt [(1-p_r_)/(n*p_r_)]* and can be compared with the measured values. Therefore, the number of spikes that are needed to reduce the variability to achieve a given CV is *n = (1-p_r_*)/(*p_r_*CV*^2^). Table 1 gives averaging time windows *T = n/R*, where *R* is spiking rate of the presynaptic axon, for representative values of *p_r_* and a range of spiking rates. Accounting for other known sources of variability at dendritic spines would require even longer time windows.

**Table 1.**
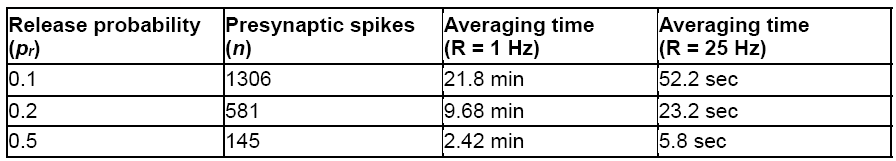
Lower bounds on time window for averaging binomially distributed synaptic input to achieve CV=0.083.

Phosphorylation of calcium/calmodulin-dependent protein kinase II (CaMKII), required for spike-timing dependent plasticity processes, integrates calcium signals over minutes to hours and is a critical step in enzyme cascades leading to structural changes induced by long-term potentiation (LTP) and long-term depression (LTD) ^29^, including rearrangements of the cytoskeleton ^30^. The time course over which CaMKII integrates calcium signals is within the range of time windows we predict would be needed for averaging (Table 1). Similar time windows occur in synaptic tagging and capture: Inputs that are too weak to trigger LTP or LTD can be “rescued” by a stronger input to neighboring synapses if it occurs within an hour ^31,32^, which also requires CaMKII ^33,34^.

Due to the many sources of variability, information encoded at a single synapse cannot be read out with a single input spike. Read out of the information over multiple spikes might reflect a sampling strategy designed for energetic efficiency since it is the physical substrate that must be stable for long-term memory retention, not the read out of individual spikes ^35^.

Previous lower bounds on the precision of synaptic strength in the hippocampus were based on whole spine volume ^9,36^. Our estimate based on spine head volume is an order of magnitude greater precision. Complementing our observations and analysis, highly correlated *p_r_* at multiple contacts between the axon of a given layer 2/3 pyramidal neuron and the same target cell has been reported ^37^. Thus, our findings suggest that 3DEM measurements of neocortical dendritic spines may reveal similarly precise estimates of synaptic efficacy.

## Acknowledgements

We are grateful to Dr. Mary Kennedy, Dr. Charles Stevens, Dr. Cian O’Donnell, and Dr. Krishnan Padmanabhan, for discussions on many aspects of synaptic spines and CaMKII, Josef Spacek, and Dylan Yokoyama for data acquisition, and Libby Perry and Robert Smith for serial sectioning and image acquisition. This research was supported by NIH grants NS44306 (to Mary Kennedy), NS21184, MH095980, and NS074644 (to Kristen Harris), P41-GM103712, MH079076 and the Howard Hughes Medical Institute (to T. Sejnowski).

## Supplementary Materials

Materials and Methods

Figures S1-S12

